# FAME: Fragment-based Conditional Molecular Generation for Phenotypic Drug Discovery

**DOI:** 10.1101/2022.01.21.477292

**Authors:** Thai-Hoang Pham, Lei Xie, Ping Zhang

## Abstract

*De novo* molecular design is a key challenge in drug discovery due to the complexity of chemical space. With the availability of molecular datasets and advances in machine learning, many deep generative models are proposed for generating novel molecules with desired properties. However, most of the existing models focus only on molecular distribution learning and target-based molecular design, thereby hindering their potentials in real-world applications. In drug discovery, phenotypic molecular design has advantages over target-based molecular design, especially in first-in-class drug discovery. In this work, we propose the first deep graph generative model (FAME) targeting phenotypic molecular design, in particular gene expression-based molecular design. FAME leverages a conditional variational autoencoder framework to learn the conditional distribution generating molecules from gene expression profiles. However, this distribution is difficult to learn due to the complexity of the molecular space and the noisy phenomenon in gene expression data. To tackle these issues, a gene expression denoising (GED) model that employs contrastive objective function is first proposed to reduce noise from gene expression data. FAME is then designed to treat molecules as the sequences of fragments and learn to generate these fragments in autoregressive manner. By leveraging this fragment-based generation strategy and the denoised gene expression profiles, FAME can generate novel molecules with a high validity rate and desired biological activity. The experimental results show that FAME outperforms existing methods including both SMILES-based and graph-based deep generative models for phenotypic molecular design. Furthermore, the effective mechanism for reducing noise in gene expression data proposed in our study can be applied to omics data modeling in general for facilitating phenotypic drug discovery.

## 1 Introduction

*De novo* molecular design which requires knowledge from multidisciplinary domains including chemistry, biology, and computational science is a challenging task in drug discovery by virtue of the complexity in the corresponding molecular space [1]. This task aims to generate novel chemical compounds with desirable pharmacological properties using computational methods. With the advances in computational technologies and theoretical findings in deep learning recently, many deep generative models have been shown to be powerful tools to capture complex patterns in molecular spaces, thereby enabling them to generate novel compounds with desired properties. In particular, these methods have been applied to generate both SMILES linearization [2, 3] and graph [4, 5, 6] representations of molecules. In spite of their success demonstrated by both theoretical and experimental evidences, they are designed only for the task of general molecular distribution learning [4, 5] and target-based molecular design [3], thereby hindering the performance of deep generative models in the real scenario of drug discovery.

Phenotypic molecular design is another approach in drug discovery that evaluates different chemicals against phenotypes which are characteristics observed in biological systems such as animals or cells. This approach has been shown to be more effective than the target-based approach in first-in-class drug discovery [7, 8]. Different from general molecular distribution learning and target-based molecular design which are formulated as unconditional generation and fine-tuning settings respectively, phenotypic molecular design can be considered as a conditional generation problem in which the conditions a.k.a phenotypes (e.g., gene expressions, cell images) are formulated as numerical representations (e.g., vectors, matrices, or tensors) (as shown in Figure 1). The availability of high-throughput drug-induced gene expression data [9, 10] creates opportunities for deep generative models to design novel chemicals with desired biological activities. However, existing deep generative architectures are not designed specifically for handling the following challenges in phenotype readouts, thereby making them not well-suited for phenotypic molecular design.

**Figure 1:**
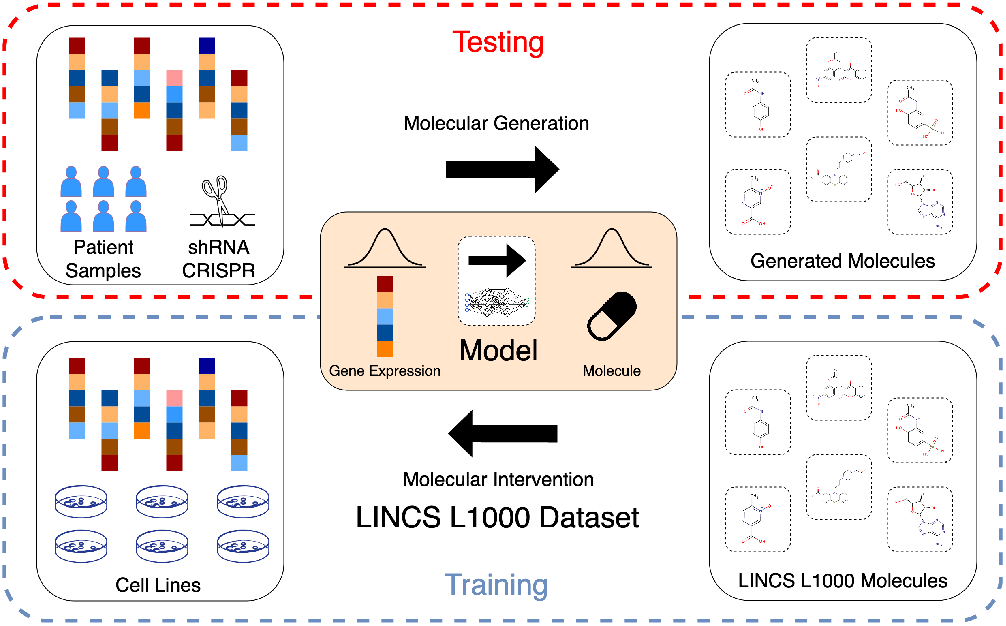
Phenotypic molecular generation. This task aims to generate novel molecules that have a high association with the corresponding phenotypes. In our setting, we train our model on LINCS L1000 dataset which consists gene expression profiles - the molecular phenotype that captures the change in expressions of multiple landmark genes in the cell line under molecular interventions. At the inference stage, given gene expression profiles retrieved from analyzing patient samples or using genetic modification techniques, we sample nove molecules that are likely to induce these profiles using our trained model.

**C1.** High-throughput phenotypic data such as drug-induced gene expression is collected in a massive, fast manner making it extremely noisy. Lacking a procedure to denoise this data precludes deep generative models from realizing their full potentials in the phenotypic molecular design.

**C2.** Existing approaches for decoding novel chemicals through their SMILES (e.g., character-by-character and context-free grammar) and graph (e.g., one-shot and node-by-node) representations are not very effective in preserving desirable pharmacological properties, making the generated molecules not associated with the corresponding phenotypes.

**C3.** Common evaluation metrics used in the general molecular distribution learning cannot provide comprehensive assessments for deep generative models in the task of generating novel chemicals that are likely to induce the phenotype of interest.

In this paper, we propose a **<u>f</u>**ragment-based condition**<u>a</u>**l **<u>m</u>**olecular g**<u>e</u>**neration model (**FAME**) for phenotypic molecular design that overcomes the afore-mentioned shortcomings in deep generative models. FAME leverages conditional variational autoencoder (VAE) to sample a latent vector from the latent space constructed during training, then combines it with the drug-induced gene expression profile which is the phenotype in our setting to generate novel molecules that are most likely to induce that desired gene expression profile. In particular, we propose the **<u>g</u>**ene **<u>e</u>**xpression **<u>d</u>**enoising model (**GED**) (dealing with **C1**) utilizing contrastive objective function to reduce noise in gene expression data by forcing gene expression profiles of the same chemical more similar than those from different chemicals. The conditional graph-based encoder of the proposed model is then used to generate the latent vector for the chemical from their molecular structure and the corresponding gene expression generated by GED. The latent vector generated from the encoder (standard normal distribution in the inference stage) and the gene expression profile are put into the fragment-based graph decoder (dealing with **C2**) to generate molecule through the sequence of fragments generated in an autoregressive manner. By leveraging the fragment-based generation strategy, the proposed model can generate novel chemicals with a high validity rate and desired biological activity. We demonstrate the effectiveness of the proposed model compared to existing deep generative models by conducting experiments on LINCS L1000 dataset [9] which consists of the measurement of differential gene expression of the most informative genes due to molecular interventions. We then evaluate the performances of these models by using Fréchet ChemNet Distance (FCD) metric [11] (dealing with **C3**) along with other common metrics used in the molecular distribution learning task. In summary, our contributions include the following:

- We design a deep generative architecture (FAME)^1^ that effectively generates novel molecules associated with the input gene expression profiles.
- We develop a gene expression denoising model (GED) utilizing contrastive objective function to successfully reduce noise in gene expression data.
- We also introduce the fragment-based graph decoder that generates novel molecules by the fragment-to-fragment strategy to achieve high validity rate and desired biological association.
- Finally, we conduct a comprehensive empirical study to demonstrate the effectiveness of FAME compared to a wide range of previous approaches for phenotypic molecular design.

## 2 Related Works

### Deep generative models in drug discovery

The abundance of molecular data generated in recent years has created an unprecedented opportunity to apply deep generative methods for molecular design. Most of these works focus on the three main tasks including molecular distribution learning, molecular optimization, and target-based molecular design. In particular, the molecular distribution learning task can be considered as unconditional generation problem in which the deep generative models are trained on the large set of molecules to learn the underlying distribution that generates these molecules. Several deep generative architecture including generative adversarial network (GAN), variational autoencoder (VAE), adversarial autoencoder (AAE), autoregressive (AR)-based and flow-based models have been proposed to learn the complex distribution of observed molecule space by considering molecules as SMILES codes (e.g., ChemVAE [2], ORGAN [12], SD-VAE [13]) or graphs of atoms and bonds (MolGAN [6], JT-VAE [5], MolecularRNN [14], MoFlow [15], GraphAF [16]). Molecular optimization is the task of produce novel molecules with optimal criteria such as octanol-water partition coefficients (logP) or drug-likeness score (QED) starting from input molecules. This task can be handled by using optimization methods (e.g., Bayesian optimization, stochastic gradient descent) to find optimum molecules on latent space [2, 5] or molecular space [17]. The target-based molecular design aims to improve a molecule’s biological activity against biological targets. This task can be handled by using a fine-tuning approach in which transfer learning [3] and reinforcement learning [18] techniques are applied to guide the deep generative models to generate novel molecules with desired biological activity.

### Phenotypic molecular design

In contrast to the target-based approach, phenotypic molecular design does not rely on knowledge of the identification of a specific molecular target to find potential drug treatments for diseases. The phenotypic approach is successful in delivering first-in-class drugs by addressing the incomplete understanding of complexity of diseases through phenotypic readouts [8]. With the advancement of cell-based phenotypic screening technologies, many drug-induced phenotypic datasets such as gene expression profiles [9] and cell painting images [10] are available for phenotypic drug discovery. These huge datasets also create opportunities for applying deep generative models for designing novel molecules with desired biological activities. However, to the best of our knowledge, only two studies focus on developing deep generative models for phenotypic molecular design [19, 20]. Both of them formulate this problem under conditional generation setting and use SMILES to represent molecules. While [19] leverages conditional GAN, [20] utilizes conditional AAE to generate novel molecules having a high association with the input phenotype.

## 3 Method

We first describe the phenotypic molecular design and notations used in our study and then introduce our deep generative model for this task including fragment-based conditional molecular generation (FAME) and gene expression denoising (GED) network. The overall architecture of FAME is shown in Figure 2.

**Figure 2:**
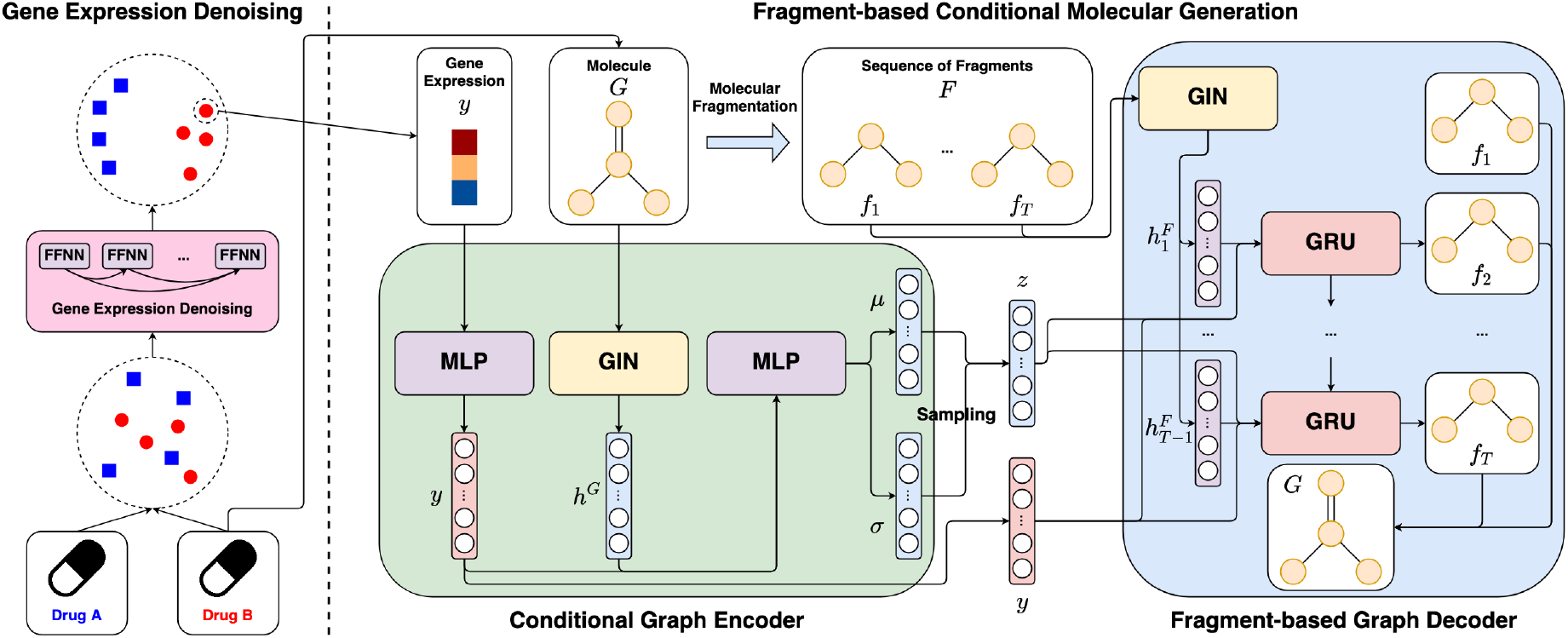
Overall architectures of GED and FAME. First, GED is used to reduce noise in gene expression profiles by mapping them to the embedding space using contrastive objective function. Then, FAME takes both gene expression embeddings and molecules as input and learns to generate the sequence of fragments constructing the input molecule. In the inference stage, the latent vector sampled from 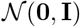 and the gene expression embedding is used to generate novel molecules that are likely to induced that embedding. (FFNN: feed-forward neural network, MLP: multi-layer perceptron, GIN: graph isomorphism network, GRU: gated recurrent unit.)

### 3.1 Problem Formulation

#### Phenotypic molecular design

We formulate this problem under conditional generation setting in which the model aims to learn a parameterized conditional distribution *P_θ_* (molecule | phenotype) and then sample novel molecules from the input phenotypes using the learned distribution. In FAME, a molecule is represented as an undirected graph *G* = (*V, E*) where *V* = {*v*_1_, *v*_2_ …, *v_N_*} and *E* = {(*v_i_*, *v_j_*)|*v_i_,v_j_* ∈ *V*} are the sets of atoms and bonds belonging to that molecule and *N* is the number of atoms. The numerical representation of graph *G* includes a node feature matrix *X* ∈ {0,1}^*N*×|*V*|^ and an adjacency tensor *A* ∈ {0,1 }^*N*×*N*×|*E*|^. A phenotype in our setting is a gene expression profile represented by a numerical vector 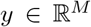 where *M* is the number of genes in that profile. Then, the goal of our proposed model is to learn a distribution *P_θ_*(*G*|*y*).

#### Conditional variational autoencoder

The conditional distribution *P_θ_* (*G*|*y*) is often intractable so stochastic gradient variational Bayes setting is applied to optimize the variational lower bound of the log-likelihood as a surrogate objective function as follows.

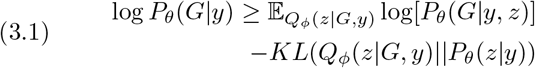

where *Q* is the learned posterior distribution (i.e., encoder network), *ϕ* and *θ* are parameters of encoder and decoder networks of the conditional VAE framework and *z* is the latent vector in the latent space constructed by this framework. The sampling process of this framework is as follows. For given gene expression profile *y*, *z* is drawn from the prior distribution 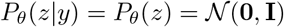 (i.e., making the latent variables statistically independent of gene expression profile), and the output graph *G* is generated from the distribution *P_θ_* (*G*|*y,z*).

### 3.2 Fragment-based Conditional Molecular Generation

The key challenge in estimating the conditional distribution is the complex space consisting of all configurations of labeled nodes and edges, which are intractable for reasonably sized graphs. We will show that existing graph-based generative models using node-by-node sampling strategy are not ideal at discovering common substructures such as rings in molecules in the experimental section, thereby making the conditional distribution hard to learn. To alleviate this phenomenon, we break molecules to a sequence of fragments and then let FAME learn to design novel molecules by sampling each fragment at each step in an autoregressive manner. This approach guarantees the model to generate valid substructures at each step by transforming the conditional distribution from the combinatorial space of atoms and bonds to the fragment space. In particular, we propose the molecular fragmentation algorithm using BRICS bonds [21] to sequentially break an input molecule to smaller substructures. The pseudo-code and example of this algorithm are shown in Algorithm 1 and Figure 3, respectively. Then, the conditional distribution is computed as follows.

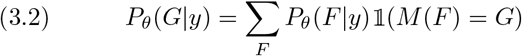

where *F* is the sequence of fragments, *M* is the reconstruction operator that maps a sequence of fragments to molecule, and 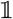 is the identity function. Note that, due to the randomness in selecting BRICS bonds in the molecular fragmentation algorithm, one molecule can have multiple fragment sequences. Under an autoregressive manner, this distribution can be further decomposed as follows.

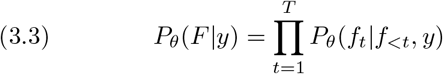

where *F* = [*f*_1_, *f*_2_, … *f_T_*] and *f*_<*t*_ = {*f*_1_, *f*_2_, … *f*_*t*−1_} Then, the objective function for FAME is as follows.

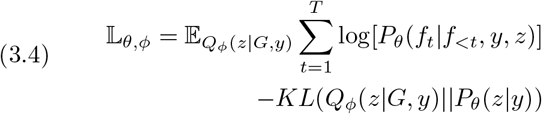

#### Algorithm 1: Molecular Fragmentation

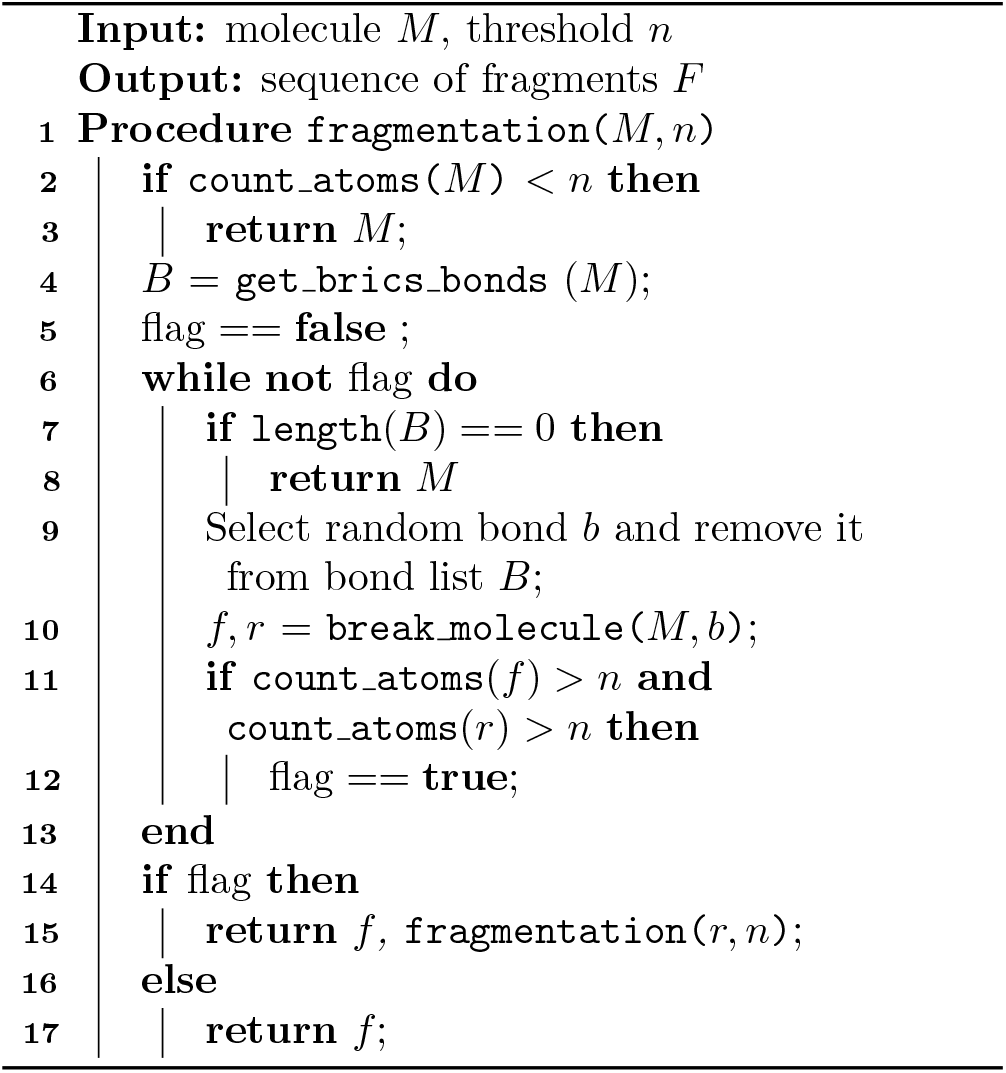

**Figure 3:**
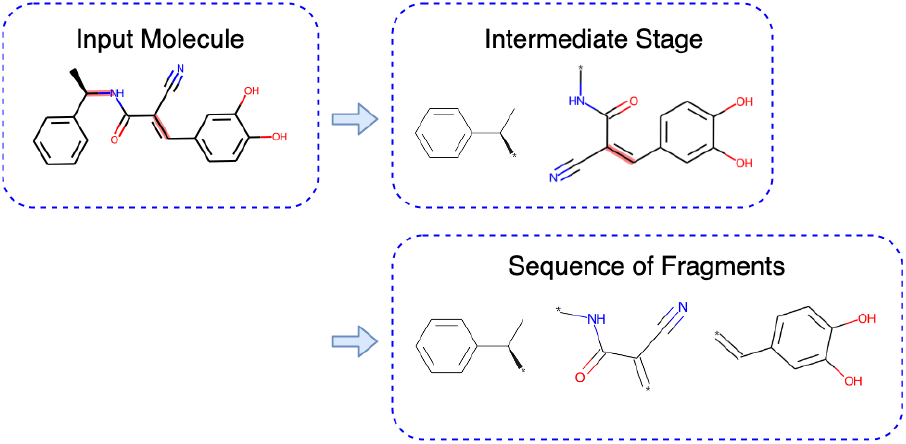
Molecular fragmentation. In this example, molecules *Flavanone* are transformed to a sequence of 3 fragments by sequentially breaking BRICS bonds (highlighted with red color) and replacing them with dummy atoms (denoted by ‘*’).

#### Conditional graph encoder

We implement the conditional graph encoder *Q_ϕ_* (*z*|*G*, *y*) as a multi-layer message passing network that leverages the graph structure of molecules and gene expression profile to construct the latent vector *z*. In particular, a 5-layer graph isomorphism network (GIN) which has been shown to be more powerful than other graph neural networks [22] is employed to learn the vector representation *h^G^* for the input graph *G* = (*X*, *A*) as follows.

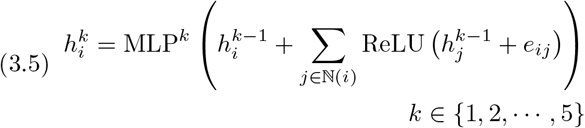

where MLP is a multi-layer feed-forward neural network, 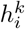 is the representation of node *v_i_* at *k* – *th* layer, 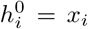 is the *k* – *th* row of the node feature matrix *X* denoting the atom type of node *v_i_*, *e_ij_* is the (*i,j*)-vector in the adjacency tensor *A* denoting the bond type of the edge between nodes *v_i_* and *v_j_*, and *N_i_* is the set of nodes that have edges to node *v_i_*. We average the representations of all nodes to generate the graph representation *h^G^* (The original model uses sum operator but we found that it makes KL-divergence loss unstable during training). Then, latent vector *z* is sampled from *N*(*μ,σ*) where *μ, σ* is computed as:

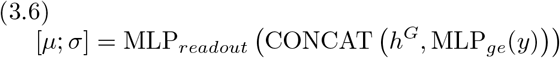

where CONCAT is the concatenation operator.

#### Fragment-based graph decoder

The latent vector *z* generated by the encoder network and the gene expression profile *y* are put into the decoder network to estimate the distribution *P_θ_* (*F*|*y, z*) which is equal to the product of conditional distributions *P_θ_*(*f_t_*|*f_<t_, y, z*). In particular, we employ GIN and gate recurrent unit (GRU) networks to model this sequence of conditional distributions as follows.

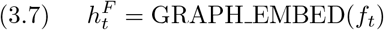

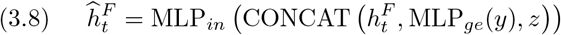

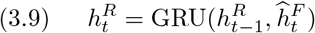

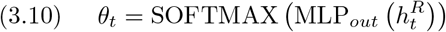

where GRAPH_EMBED is the GIN used to transform fragment *F_t_* to the vector representation 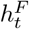, and has similar architecture to the one used in the conditional graph encoder. 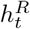 is the hidden representation computed by GRU at time step *t* and *θ_t_* is the vector that represents the conditional distribution *P_θ_*(*f_t_*|*f_<t_*, *y*, *z*).

### 3.3 Gene Expression Denoising

With the advancement of phenotypic screening technologies, several massive phenotypic datasets have been generated in a quick manner but these screening methods also introduce lots of noises in their measurements. In particular, the gene expression profiles in LINCS L1000 dataset measured under the same condition (i.e., chemical and cell line) may be very different, resulting in difficulties for deep generative models to learn the relationship between chemical and its biological activity. To alleviate this problem, we propose a gene expression denoising (GED) network utilizing contrastive objective function to map gene expression profiles into a unit hyper-sphere space and then forcing to pull together the gene expression embeddings of the same chemicals while simultaneously pushing them away from gene expression embeddings of other chemicals in that space, thereby helping to denoise gene expression data. In particular, a gene expression profile *y_l_* induced by chemical *c_l_* is projected to unit hyper-sphere space as 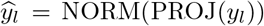 where PROJ is the projection network and NORM is the normalization operator making the learned representation to lie in the embedding space. Then the contrastive objective function is applied as follows.

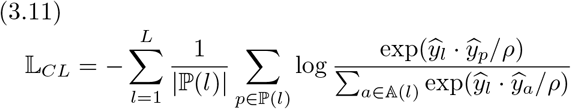

where 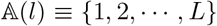 \ {*l*} is the set of all indices except *l* and 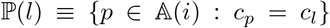 is the set of indices of all gene expression profiles induced by chemical *c_l_*. Inspired by the success of DenseNet used in image recognition task [23], we design the PROJ network as a very deep feed-forward neural network having shortcut connections between every layers as follows.

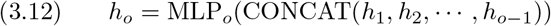

where input at *o* – *th* layer is the concatenation of all hidden representations produced in previous layers. In particular, GED consists of 64 feed-forward layers with growth rate = 16, and the size of the output layer is set at 64. After training, we replace *y_l_* by 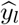 transforming the condition distribution in Equation 3.1 to 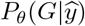.

### 3.4 Handling Infrequent Fragments

Due to the complexity of molecular space, a small number of fragments appears much more frequent than the others in the data, resulting in difficulties in estimating the probability of generating infrequent fragments. To ease this problem, at the training stage, we group all infrequent fragments and mark them with the special tag < *RARE* >. At the time step *t* of the decoder network, if *f*_*t*+1_ is infrequent fragment, the label for this step is the tag < *RARE* > instead of *f*_*t*+1_. At the inference stage, for each gene expression profile 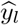 in the test set, we construct its neighboring gene expression set by calculating similarity scores between this gene expression profile and all profiles in the training set as 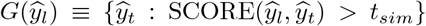 where SCORE function is Pearson’s correlation and *t_sim_* is the threshold. Then, the infrequent fragments of molecules corresponding to the gene expression profiles in this neighboring set are extracted to form the set 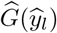. At the time step *t* of the molecular sampling process for gene expression profile 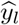, if the tag < *RARE* > is selected, the fragment *f*_*t*+1_ will be uniformly sampled from set 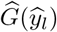 as the output of that time step.

## 4 Experiments and Discussions

In this section, we evaluate the performance of FAME on the drug-induced gene expression data and compare its results with state-of-the-art deep generative models including both SMILES-based and graph-based models designed for either unconditional or conditional generation settings to demonstrate the efficiency of our method for phenotypic molecular design. Besides achieving a superior generation performance, we also show the effectiveness of GED model in the task of reducing noise for gene expression data.

### 4.1 Datasets

The datasets used in our study include LINCS L1000, ExCAPE, and ChEMBL. The summarized statistics of these datasets are shown in Tables 1 and 2 and their details are described as follows.

**Table 1:** LINCS L1000 and ChEMBL data statistics. (ge: gene expression, mol: molecule, avg: average)

| Datasets |  | #ge | #unique<br>mols | #avg atoms<br>per mol | #atom<br>types |
| --- | --- | --- | --- | --- | --- |
| LINCS L1000 | Training | 25658 | 16019 | 31.23 | 15 |
|  | Validation | 2872 | 1782 | 30.83 |  |
|  | Testing | 3291 | 1980 | 30.54 |  |
| ChEMBL | Training | n/a | 1290143 | 30.13 | 33 |
|  | Validation | n/a | 184488 | 30.12 |  |
|  | Testing | n/a | 368603 | 30.12 |  |

**Table 2:**
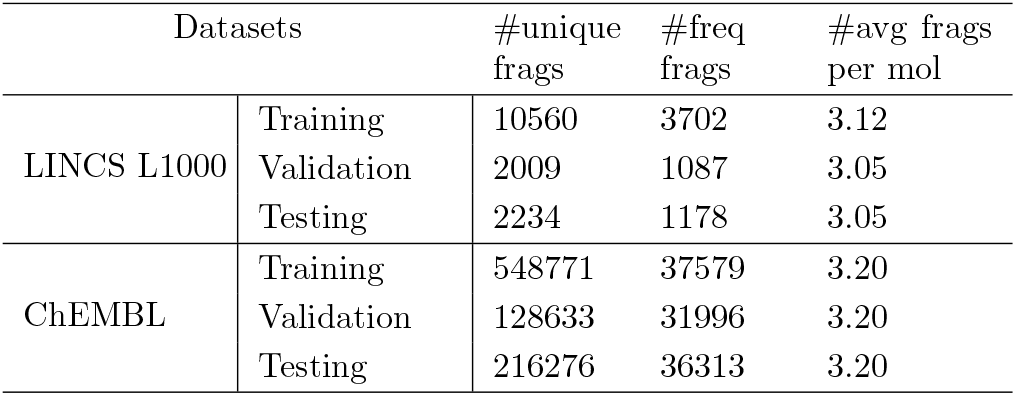
Fragment statistics in LINCS L1000 and ChEMBL datasets. (frag: fragment, freq: frequent, mol: molecule, avg: average)

#### LINCS L1000 dataset

This dataset consists of the measurements of gene expression changes for most informative genes (i.e., 978 landmark genes) caused by molecular interventions at a variety of time points, doses, and cell lines [9]. The dataset has 5 levels. We use the level 4 (gene expression profiles measured for each bio-replicate) and the level 5 (gene expression profiles which are averaged from profiles of corresponding bioreplicates). In our study, we conduct experiments on the subset of this dataset extracted by [19] which consists 31,821 level 5 gene expression profiles induced by 19, 768 compounds in MCF7 and VCAP cell lines with the largest doses (i.e., 5 and 10 *μM*) after 24 hours of exposure. These gene expression profiles are considered as chemical-induced phenotypes and are used to train the generative model. We split this dataset into training, validation, and testing set with a ratio 80 : 10 : 10 in terms of molecules. The testing set is referred to as the internal testing set in our study. We also use level 4 gene expression profiles to train GED model to reduce noises in the gene expression data.

#### ExCAPE dataset

Each gene expression profile has only one corresponding molecule as a reference, thereby prohibiting the usage of statistical metrics to evaluate the performance of generative models for molecular generation task at the individual profile level. To surpass this problem, we further evaluate the performances of generative models on the dataset constructed in [19]. In particular, 148 gene expression profiles caused by the intervention of CRISPR technology to the ten protein targets (i.e., SMAD3, TP53, EGFR, AKT1, AURKB, CTSK, MTOR, AKT2, PIK3CA, HDAC1) are selected as phenotypes, and then the known active molecules for these targets are extracted from the ExCAPE dataset [25] to serve as the reference molecule sets for each of these targets (i.e., > 1,000 molecules for each target/gene expression profile). We refer to this dataset as the external testing set in our study.

#### ChEMBL dataset

Deep generative model often requires large training data to achieve good performances while the size of LINCS L1000 dataset is relatively small in terms of the number of molecules. Thus, we utilize the ChEMBL dataset [24] which consists of ~ 2,000,000 drug-like molecules to pre-train the deep generative models for the distribution learning task, and then helping them to recognize important patterns in the molecular space.

### 4.2 Experimental Settings

#### Baseline models

To validate the performance of FAME for the phenotypic molecular design, we compare it with a wide range of deep generative models including both SMILES-based and graph-based models. To the best of our knowledge, there are only two SMILES-based models designed specifically for phenotypic molecular design [19, 20]. Thus, we adapt some graph-based models for this task by incorporating gene expression profiles into these models. The details of these models are presented as follows.

- **UniAAE** [20]. This SMILES-based model leverages conditional AAE framework to explicitly learn the shared and separated latent representations for molecules and gene expression profiles, and then the shared representations of gene expression profiles are used to sample novel molecules.
- **LatentGAN** [19]. This SMILES-based model leverages conditional Wasserstein GAN with a gradient penalty to generate latent representations for molecules from the input gene expression profiles. Then, the pre-trained auto-encoder is used to decode these latent vectors for novel molecules.
- **MolGAN** [6]. The graph-based model leverages GAN architecture to generate node feature matrix and adjacency tensor from noise vector. To make it work in the conditional generation setting, the generator takes input as the concatenation of gene expression profiles and noise vectors. We apply the post-processing technique proposed in [15] to guarantee the chemical validity of generated graphs.
- **MolecularRNN.** [14] This graph-based autoregressive model handles molecular graphs by incorporating atom and bond labels into its architecture. This model guarantees the chemical validity by using valency-based rejection sampling method. We concatenate the gene expression profile with inputs of NodeRNN component to adapt this model into the conditional generation setting.

#### Evaluation metrics

The most important criterion of phenotypic molecular design is to generate novel molecules with desired biological activity (i.e., molecules that are likely to induce the input gene expression profiles). Thus, we focus on measuring the similarity between reference and generated molecules by utilizing Fréchet ChemNet Distance (**FCD**) [11]. This metrics is computed from hidden representations of molecules generated by the model trained to predict drug activities so it can measure the similarity w.r.t both chemical and biological perspectives. Besides FCD, we also report results for widely-used metrics that measure the general quality of generated molecules such as **Valid** (the chemical validity rate of generated molecules), **Novel** (the fraction of generated valid molecules which are not in the training dataset), **Unique** (the fraction of unique correct molecules), and **Internal Diversity** (the average distance between generated molecules).

## 4.3 Results

We conduct experiments to answer the following questions.

- **Q1.** How effective is FAME for phenotypic molecular design compared with previous works?
- **Q2.** How well does GED reduce noise in gene expression data by contrastive loss function?

### Phenotypic Molecular Design

To evaluate whether generated molecules can induce the input gene expression profiles, we compare it with the set of reference molecules by using FCD metric to calculate the distance with respect to chemical and biological perspectives between these two sets. For the internal testing set, we calculate the FCD metric between generated set and the whole testing set while for the external testing set, we calculate the FCD metric between generated and reference sets of each gene expression profile and report the average result over these profiles. As shown in Table 3, FAME achieves the smallest distances (i.e., FCD scores) at both internal and external testing sets making its generated molecules to be most similar to the reference molecules compared to other models. This result shows the effectiveness of using graph representation, fragment-to-fragment sampling strategy, and gene expression denoising model to estimate the conditional distribution in Equation 3.1. For other metrics, performances of FAME are on par with other deep generative models, thereby showing the comprehensiveness of the proposed model.

**Table 3:**
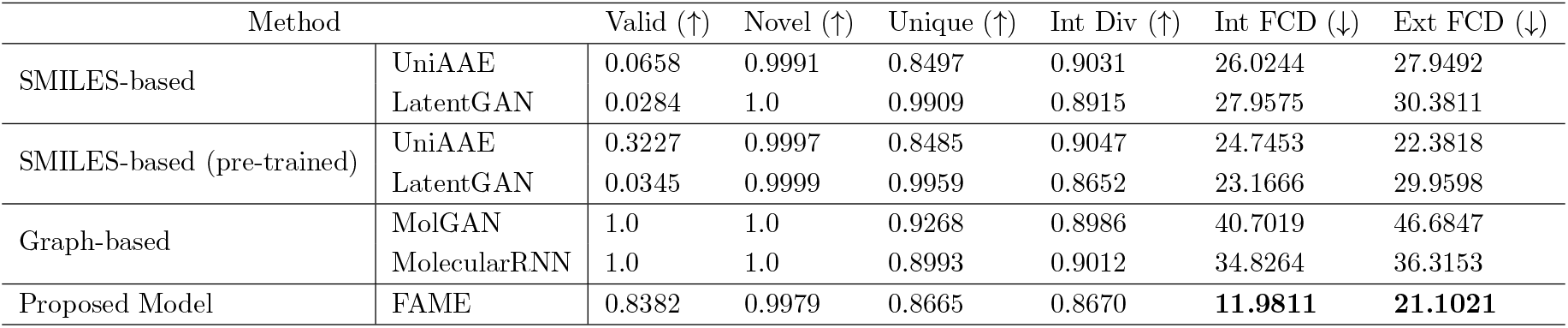
Performances of FAME and baseline models for phenotypic molecular design. The directions of the arrows indicate the optimized directions of the metrics. (Int Dive: internal diversity, Int FCD: FCD measured on internal test set, Ext FCD: FCD measured on external test set).

For baseline models, we observe that SMILES-based models achieve significantly better performances compared to the graph-based models in terms of FCD. To investigate this phenomenon, we visualize the generated molecules of these models in Figure 4. We can see that graph-based models can easily exploit common metrics by using post-processing methods to make the generated molecules chemically valid. However, these methods cannot guarantee the generated molecules to preserve substructures such as rings, thereby making them not drug-like molecules. As shown in Figure 4, MolGAN and MolecularRNN often generate molecules with incomplete rings or infrequent substructures. For SMILES-based models, these substructures can be easily recognized because they are often substrings in the SMILES code (e.g., ‘c1ccccc1’ is the substructure/ring of ‘NC1C[C@H]1c1ccccc1’) but these models have low validity rates. Our proposed model combines the strengths of these two approaches resulting in a high validity rate and substructure preservation.

**Figure 4:**
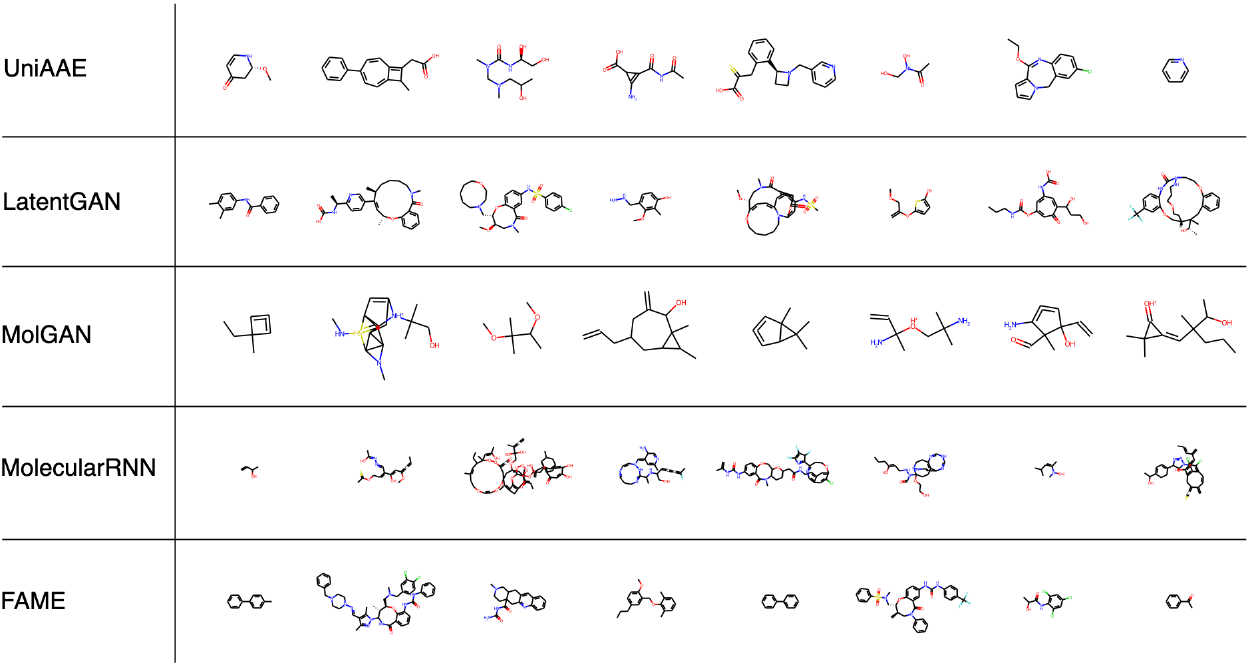
Sample molecules generated by UniAAE, LatentGAN, MolGAN, MolecularRNN, and FAME.

### Gene Expression Denoising

To investigate the contribution of GED model to the phenotypic molecular design, we compare the performances of UniAAE and LatentGAN using the original gene expression profiles with those using the gene expression embeddings generated by GED. The metrics used in this experiment are negative log-likelihood (NLL) and FCD. As shown in Table 4, using denoised gene expression embeddings improves the performances of these two generative models. Specifically, both of these models have lower NLL (fit the learned distribution better) and FCD (generate molecules having stronger associations with gene expression profiles) scores when incorporating GED.

**Table 4:**
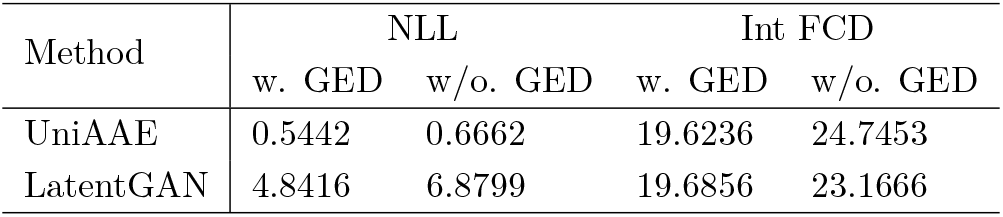
Performances of UniAAE and LatentGAN with and without incporporating GED.

| Method | NLL |  | Int FCD |  |
| --- | --- | --- | --- | --- |
|  | w. GED | w/o. GED | w. GED | w/o. GED |
| UniAAE | 0.5442 | 0.6662 | 19.6236 | 24.7453 |
| LatentGAN | 4.8416 | 6.8799 | 19.6856 | 23.1666 |

## 5 Conclusion

Phenotypic molecular design is a crucial problem in drug discovery. In this paper, we propose a fragment-based conditional molecular generation model (FAME) for this task by formulating it under a conditional VAE framework to learn the conditional distribution that generates molecules from gene expression profiles. To tackle the issues of learning this complex distribution, FAME transforms this distribution from combinatorial space of atoms and bonds to a fragment space and then learns to generate the sequence of fragments in an autoregressive manner. Moreover, the gene expression denoising (GED) model is proposed to handling noises in gene expression data by leveraging a contrastive objective function. The experimental results demonstrate that our proposed model outperforms other state-of-the-art deep generative models for phenotypic molecular design. Moreover, the denoising mechanism proposed in our study could is valuable addition to be applied to other phenotypic drug discovery applications using gene expression data.

## Acknowledgments

This work was supported in part by research grants from NIH (NIGMS R01GM141279) and OSU BMI Pilot Grant Award.

## Footnotes

1 Code and data are available at https://github.com/pthl993/FAME

